# Into new depths: climate-driven habitat expansion of the endangered skate *Dipturus chilensis* (Chondrichthyes, Rajiformes)

**DOI:** 10.64898/2026.03.26.714520

**Authors:** Jaime A. Villafaña, Diego Almendras, Daniel González-Aragón, Felipe I. Torres, Francisco J. Concha, Ana B. Guzmán-Castellanos, Ignacio Contreras, Karina E. Buldrini, Pablo Oyanadel-Urbina, Carolina A. Sandoval, Benjamín Miranda, Gabriel Mazo, Felipe Cárdenas, Mathias Valdivia, German Pequeño, Carlos Lara, Marcelo M. Rivadeneira

**Affiliations:** Departamento de Ecología, Facultad de Ciencias, Universidad Católica de La Santísima Concepción, Concepción, Chile; Proyecto Raya Águila, La Serena, Chile; Programa de Doctorado en Ciencias Mención Biodiversidad y Biorecursos, Universidad Católica de la Santísima Concepción, Concepción, Chile; Instituto Milenio en Socio-Ecología Costera (SECOS), Santiago, Chile; Laboratorio de Biología y Conservación de Condrictios (Chondrolab), Facultad de Ciencias del Mar y de Recursos Naturales, Universidad de Valparaíso, Chile; Departamento de Ciencias Ecológicas, Facultad de Ciencias, Universidad de Chile, Chile; Área de Paleontología, Museo Nacional de Historia Natural, Santiago, Chile; Área de investigación y desarrollo, Therium Servicios Profesionales Limitada, Paleontología y Patrimonio, Chile; Laboratorio de Paleobiología, Centro de Estudios Avanzados en Zonas Áridas, Coquimbo, Chile; Instituto de Ciencias Marinas y Limnológicas, Universidad Austral de Chile, Valdivia, Chile; Centro de Investigación en Recursos Naturales y Sustentabilidad, Universidad Bernardo O’Higgins, Santiago, Chile; Departamento de Biología, Universidad de La Serena, La Serena, Chile

**Keywords:** Climate, models, ecology, elasmobranchs, Pacific Ocean

## Abstract

The yellownose skate (*Dipturus chilensis*) is an endangered skate with a narrow distribution in the southeastern Pacific, facing intense fishing pressure and potential climate threats. Using a species distribution model, we projected the current and future distribution of *D. chilensis* under contrasting climate change scenarios (SSP1-2.6, SSP2-4.5, and SSP5-8.5) for mid-century (2050) and end-of-century (2100). Our models, which demonstrated robust predictive performance significantly better than random expectations, identified maximum temperature and minimum oxygen as the primary environmental drivers of habitat suitability. Projections revealed a consistent poleward range shift towards the Channels and Fjords of Southern Chile ecoregion across all scenarios. While localized habitat loss was projected in Central Chile and Araucanian ecoregions, particularly under high emissions (SSP5-8.5), these losses were outweighed by southern expansions, leading to a net increase in total suitable habitat by 2100. These findings underscore the critical need for climate-adaptive management strategies, including the protection of emerging southern refugia and dynamic fisheries regulations, to ensure the long-term persistence of *D. chilensis*.

## 1. Introduction

The yellownose skate *Dipturus chilensis* (Guichenot, 1848), is a medium-sized, demersal hardnose skate endemic to the southeastern Pacific Ocean. Taxonomically, this species has undergone several revisions (see Concha et al., 2019). Originally placed in *Raja* Linnaeus, 1758, it was reassigned to *Dipturus* Rafinesque, 1810 by McEachran and Dunn (1998) and later moved to *Zearaja* Whitley, 1939 by Last and Gledhill (2007). However, molecular studies (Naylor et al., 2012a; Vargas-Caro et al., 2017) revealed that *Zearaja* forms a small lineage deeply nested within *Dipturus* and recognizing it as a separate genus would render *Dipturus* paraphyletic. Thus, following Concha et al. (2019), we use *Dipturus* to maintain taxonomic stability.

This species occurs along the Chilean continental shelf and upper slope from approximately 27°S to 56°S (Bustamante et al., 2014; ; Concha et al., 2019; Navia et al., 2025), inhabiting soft-bottom substrates at depths between 9 and 512 m and temperatures ranging between 7.7 and 11.5 °C, as reported within a limited temporal and spatial context by Ahumada et al. (2025) . Its life-history traits, such as late maturity, low fecundity, and high longevity, make this species highly susceptible to overexploitation (Licandeo et al., 2006; Aversa et al., 2011; Bustamante et al., 2012; Concha et al., 2012). Historically, it has been one of the most exploited elasmobranchs in Chilean waters, targeted by both small and large-scale fisheries (Quiroz et al., 2008, 2011; Wiff et al., 2018; Becerril-García et al., 2022). Landings peaked at approximately 5000 tons in 2003 but have since declined to only 537 tons in 2024 (SERNAPESCA, 2024). Currently, only the artisanal fleet targets *D. chilensis*, operating with annual quotas since 2016 and having undergone complete bans in 2014, 2015 and 2017 (San Juan et al., 2025). Additionally, it occurs as bycatch in several deep-water trawl, gillnet and longline fisheries, both artisanal and industrial (Ahumada et al., 2025; Bernal et al., 2025a, 2025b). Due to intense fishing pressure, restricted distribution and vulnerable life-history traits, the species is now classified as Endangered on the IUCN Red List of Threatened Species (Dulvy et al., 2021a).

Climate change poses additional threats to sharks and rays worldwide, compounding the impacts of overfishing through ocean warming, acidification, deoxygenation, and increased storm intensity (Dulvy et al., 2014, 2021; Gianelli et al., 2023). Those stressors have also been assessed for chondrichthyans from the Eastern Tropical Pacific (ETP) (Cerutti-Pereyra et al., 2024). Chondrichthyans are particularly sensitive to thermal stress, which can drive range shifts, elevate metabolic demands, disrupt reproduction, and alter prey availability, ultimately reducing fitness and increasing extinction risk (Simpfendorfer and Kyne, 2009; Di Santo et al., 2015; Schwieterman et al., 2019; Coelho et al., 2021, 2022; Pollom et al., 2024). Intense temperature fluctuations and progressive warming compromise embryonic development and survival in oviparous elasmobranchs (Di Santo, 2015; Ripley et al., 2021).

For instance, evidence from the Southwest Indian Ocean highlights elevated extinction risk of endemic skates under the combined pressures of climate change and overfishing (Pollom et al., 2024). Recent assessments have further documented the climate vulnerability of chondrichthyans in the Eastern Tropical Pacific, particularly deep-water species and batoids (Cerutti et al., 2024), reinforcing concerns about the sensitivity of these taxa to ocean warming In temperate and subpolar regions, thermal habitat shifts may trigger range redistributions, reshape biogeographic patterns, and increase local extinction risk for thermally sensitive species (Nye et al., 2009; Coelho et al., 2022). In this context, understanding and forecasting climate-driven range shifts require analytical tools that can explicitly link species’ occurrences to environmental variability across space and time.

Species Distribution Models (SDMs) have become essential tools in marine conservation biology for predicting climate-driven changes in species ranges, habitat loss, and community restructuring (Robinson et al., 2017; Klaassen et al., 2025). By integrating occurrence data with environmental variables, SDMs estimate species–environment relationships under current and future scenarios using statistical or theoretical approaches (Jones et al., 2012; O’Brien et al., 2022; Sillero et al., 2023). These spatially explicit predictions inform conservation planning by identifying vulnerabilities and priority areas for intervention (Karp et al., 2025). This approach is particularly relevant for skates, which exhibit high sensitivity to anthropogenic pressures (Dulvy et al., 2003; Frisk et al., 2005; Dulvy et al., 2014), and recent studies confirm the utility of SDMs in addressing key threats to these species (Klippel et al., 2016; Pennino et al., 2019; Bisch et al., 2022., Silva et al., 2023).

Despite its importance, climate change impacts on cartilaginous fishes remain a low research priority in Latin America (Becerril-García et al., 2022). Paleontological evidence suggests that past climatic fluctuations drove latitudinal shifts and extinctions in Chilean chondrichthyans (Villafaña and Rivadeneira, 2014, 2018). Contemporary studies show that elasmobranch distributions are strongly influenced by environmental conditions such as temperature, oxygen availability, and seasonal oceanographic variability, which can drive habitat shifts and redistribution across spatial scales (Papastamatiou et al., 2015; Santos et al., 2021; Bouyoucoset al., 2022; Vilmar and Di Santo, 2022). Batoids and other elasmobranchs exhibit seasonal or environmentally driven movements that allow them to track suitable thermal and ecological conditions or avoid unfavorable environments (Osgood et al., 2020; Sherman et al., 2023; Barrowclift et al., 2023). Although recent studies have explored broad-scale vulnerability patterns of chondrichthyans in the Eastern Pacific (Cerutti-Pereyra et al., 2024; Navia et al., 2025), assessments focused specifically on demersal temperate skates remain scarce. This gap underscores the need for integrated evaluations to support sustainable management and conservation of Chilean marine ecosystems.

Here, we assess the potential impacts of climate change on the distribution of the yellownose skate by the year 2100. Using a series of SDMs, we project changes in its area of occupancy under future environmental scenarios, providing insights to guide conservation and fisheries management strategies in Chile. This study represents the first climate-driven habitat modeling effort for an endangered skate in the southeastern Pacific, offering critical information for proactive conservation planning.

## 2. Methods

### 2.1 Occurrence data

We compiled a dataset with 2826 records of *D*. *chilensis* from multiple sources, including Global Biodiversity Information Facility (GBIF.org, 2024; n= 977), personal observations (n= 79), institutional online repositories (n= 16), exploratory scientific fisheries (n= 1378), and scientific literature (n=376) **(Table S1)**. We restricted our dataset to Pacific records, as recent taxonomic revisions indicate that Atlantic occurrences previously attributed to this species correspond to *D. brevicaudatus* (Concha et al., 2019). We applied spatial thinning using the ‘spThin’ R package with a distance threshold of 11 km (twice the ∼5.5 km pixel size of our environmental layers) to reduce the strong sampling bias in central and central-southern Chile, ensuring a more even spatial distribution of occurrence records for *D*. *chilensis.* After data cleaning, the final occurrences of *D*. *chilensis* along the Southeast Pacific were concentrated in Araucarian and Chiloense ecoregions, with fewer records in the Central Chile ecoregion and scattered records in the Channels and Fjords of Southern Chile ecoregion (**Fig. 1**). We applied a custom script to further correct sampling bias, which involved generating a bias file by applying kernel density estimation (KDE) to the occurrence coordinates to create a raster of sampling effort. This raster was resampled to match the resolution of the environmental layers and normalized to a 0–1 scale, producing a bias layer used in Maxent to down-weight the influence of oversampled regions.

**Figure 1.**
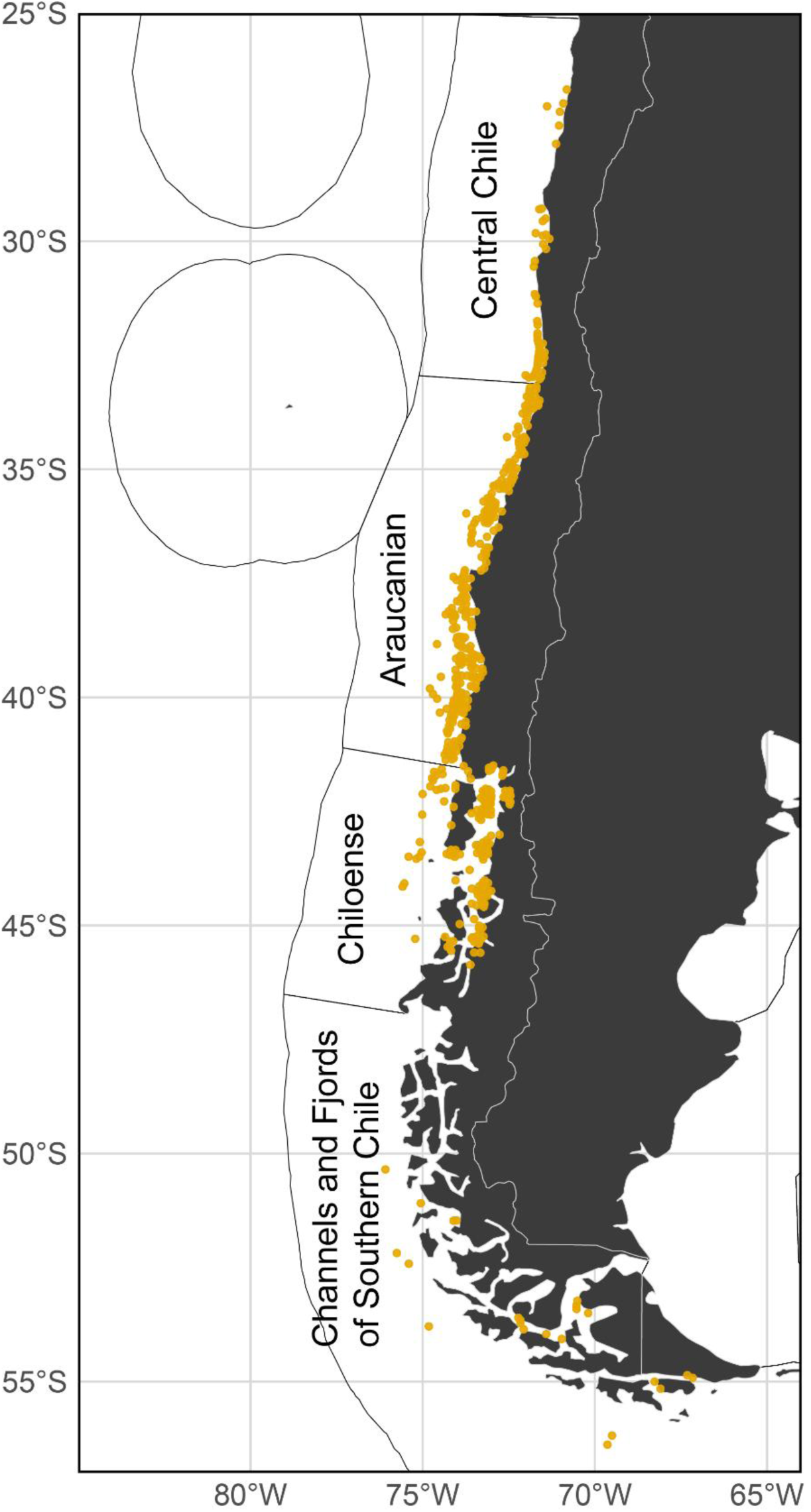
Occurrence records of *Dipturus chilensis* used in this study across the four ecoregions it inhabits: Central Chile, Araucanian, Chiloense, and Channels and Fjords of Southern Chile (Spalding et al. 2007). Records were cleaned and spatially filtered by applying a 5.5 km² grid, retaining a single randomly selected occurrence per cell to reduce spatial clustering.

### 2.2 Environmental predictors

Marine environmental variables were obtained from Bio-ORACLE version 3.0 (https://www.bio-oracle.org; Assis et al. 2024), which provides data for present and future conditions based on CMIP6 Shared Socioeconomic Pathway (SSP) scenarios from the Intergovernmental Panel on Climate Change (IPCC). These layers have a spatial resolution of 0.05° (approximately 5.5 km at the Equator). To ensure consistency across time periods, we used the same set of four environmental variables applied in the present-day model for projecting future habitat suitability for *D. chilensis*. Projections were made for two future decadal periods (2040–2050 and 2090–2100) under three contrasting greenhouse gas emission scenarios: SSP1-2.6 (“sustainable development”), SSP2-4.5 (“intermediate pathway”), and SSP5-8.5 (“fossil-fueled development”).

*D. chilensis* inhabits soft-bottom habitats across a broad depth range (9–512 m) (Ahumada et al., 2025). Therefore, all environmental predictors were extracted from the benthic layers of Bio-ORACLE (https://www.bio-oracle.org; Assis et al. 2024). The environmental predictors initially considered to construct the species distribution model included maximum and minimum temperature (°C); minimum and maximum dissolved oxygen (mmol m⁻³); maximum and minimum salinity (PSU); maximum and minimum pH; maximum and minimum sea water velocity (m.s^⁻1^); and maximum and minimum phytoplankton concentration (mg m⁻³).

A preliminary Maxent model was first generated using all candidate environmental predictors to evaluate their individual contribution. To reduce redundancy among variables, we assessed collinearity using both the Variance Inflation Factor (VIF) and Spearman’s correlation coefficient (|rₛ| > 0.75 and VIF > 5 considered high collinearity) (Dormann et al., 2013). The final selection of predictors was based on their contribution in the preliminary model, collinearity diagnostics, and ecological relevance for *D. chilensis* as reported in the literature (Licandeo et al., 2006; Bustamante et al., 2021; Ahumada et al., 2025). Accordingly, the variables retained for the final model were minimum dissolved oxygen, maximum salinity, and minimum and maximum temperature.

Predictor layers were clipped to the relevant marine biogeographic regions to define the study area where *D. chilensis* has been recorded. Restricting the analysis to these boundaries ensures the inclusion of areas accessible through natural dispersal and improves model evaluation (Sillero et al., 2021). Delimitation of biogeographic areas delineate these regions, we adopted the global classification of Marine Ecoregions of the World by Spalding et al., (2007), retaining four ecoregions that encompass the known distribution of *D. chilensis*: Central Chile, Araucanian, Chiloense, and Channels and Fjords of Southern Chile (**Fig. 1**). A mask was created based on these ecoregions, and all environmental variables were clipped accordingly to restrict the model to ecologically and biogeographically relevant areas.

### 2.3 Model performance, evaluation, threshold, and projections

We used Ecological Niche Modeling (ENM) to estimate habitat distribution of *D. chilensis* with Maxent version 3.4.4, a machine learning algorithm based on the principle of maximum entropy. This method predicts species habitat distributions from presence-only data in combination with environmental variables, generating maps of habitat suitability across the study area (Phillips and Dudík, 2008; Phillips et al., 2021). Models were run with 10 replicates using cross-validation, a maximum of 1000 iterations and 10000 background points randomly sampled from within the study extent (Phillips et al., 2006). For each fold, presence data were split into 70% for training and 30% for evaluation.

Model performance was assessed using the Area Under the Curve (AUC) of the Receiver Operating Characteristic (ROC), which measures the model ability to discriminate between presences and background conditions (Peterson et al., 2008). AUC values near 0.5 indicate random performance, whereas values closer to 1.0 reflect high predictive accuracy (Swets, 1988). This metric accounts for both sensitivity and specificity, providing a comprehensive measure of model fit (Phillips et al., 2004). To evaluate whether model performance was better than random, we built models by generating random occurrence sets of equal size to the empirical records, using identical Maxent settings. Empirical AUCs were then compared against the distribution of null AUCs. Statistical differences were tested for normality with the Shapiro–Wilk test and mean and median comparisons were performed with Welch’s t-test and the Wilcoxon rank-sum test, respectively, following Raes and ter Steege (2007).

Changes in suitable habitat were quantified across all model replicates. To convert the continuous Maxent outputs into binary maps, we applied the “Maximum training sensitivity plus specificity” (MTSS) thresholding criterion, which identifies the value that maximizes the balance between omission and commission errors and corresponds to the point of greatest combined sensitivity and specificity along the ROC curve (Liu et al., 2013). This approach is widely recommended for presence-only data as it is equivalent to maximizing the true skill statistic (TSS), providing a conservative and ecologically meaningful cutoff for distinguishing suitable habitat. For our final ensemble, the MTSS value was 0.3655, and pixels with suitability ≥ 0.3655 were classified as suitable habitat.

To quantify changes in habitat extent, all binary rasters were reprojected to an equal-area coordinate reference system (World Equidistant Cylindrical, EPSG:6933), ensuring accurate comparisons of surface areas expressed in square kilometers. Area estimates extracted from each replicate and climate scenario were compiled into a single dataset for downstream statistical analyses. All processing steps were conducted in R version 4.5.0 using the packages *terra*, *sf*, and *dplyr*.

## 3. Results

Model performance was significantly higher than expected by chance. The mean test AUC of the empirical models was 0.94, substantially higher than the mean of the null models (0.51 ± 0.04) (**Table S2**). A Wilcoxon test confirmed that empirical AUC values were significantly higher than the null distribution (W = 0, p < 0.0001), with 100% of null replicates performing below the empirical mean. These results demonstrate that the predictive ability of the empirical models cannot be explained by random processes. Among the four final predictors retained in the model, maximum temperature at depth showed the highest contribution (**Table 1**). Minimum dissolved oxygen at depth was the next most important variable, explaining a substantial portion of the variance, followed by minimum temperature at depth, while maximum salinity at depth contributed only marginally.

**Table 1.**
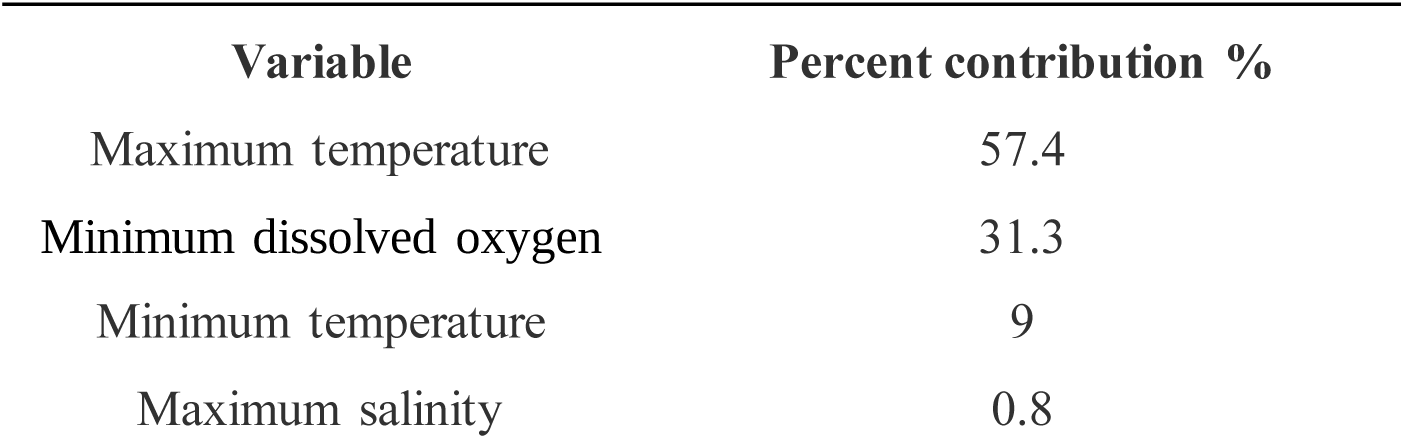
Contribution of the variables to the Maxent model of the present model of *Dipturus chilensis*. All variables correspond to depth and were downloaded from Bio-ORACLE version 3.0 (https://www.bio-oracle.org; Assis et al. 2024)

| Variable | Percent contribution % |
| --- | --- |
| Maximum temperature | 57.4 |
| Minimum dissolved oxygen | 31.3 |
| Minimum temperature | 9 |
| Maximum salinity | 0.8 |

Model projections revealed broadly consistent patterns across scenarios and time periods, characterized by a general southward range shift toward the Channels and Fjords of Southern Chile ecoregion (**Fig. 2**). For the near-future period (2040–2050), differences among the three scenarios were minimal, and most of the current suitable habitat was conserved, with only some localized gains in the Channels and Fjords of Southern Chile ecoregion. By the late century (2090–2100), clearer divergences emerged. Under SSP1-2.6, the poleward expansion was accompanied by a slight loss of habitat suitability along the northern Chilean coast (i.e., Central Chile ecoregion). Under SSP2-4.5, projected gains in the Channels and Fjords of Southern Chile ecoregion were more substantial, and little to no habitat loss was observed across the Southeast Pacific. The SSP5-8.5 scenario projected the most pronounced changes, with the largest gains in higher latitudes but also the most extensive losses, particularly along the Central Chile and Araucanian ecoregions of Chile.

**Figure 2.**
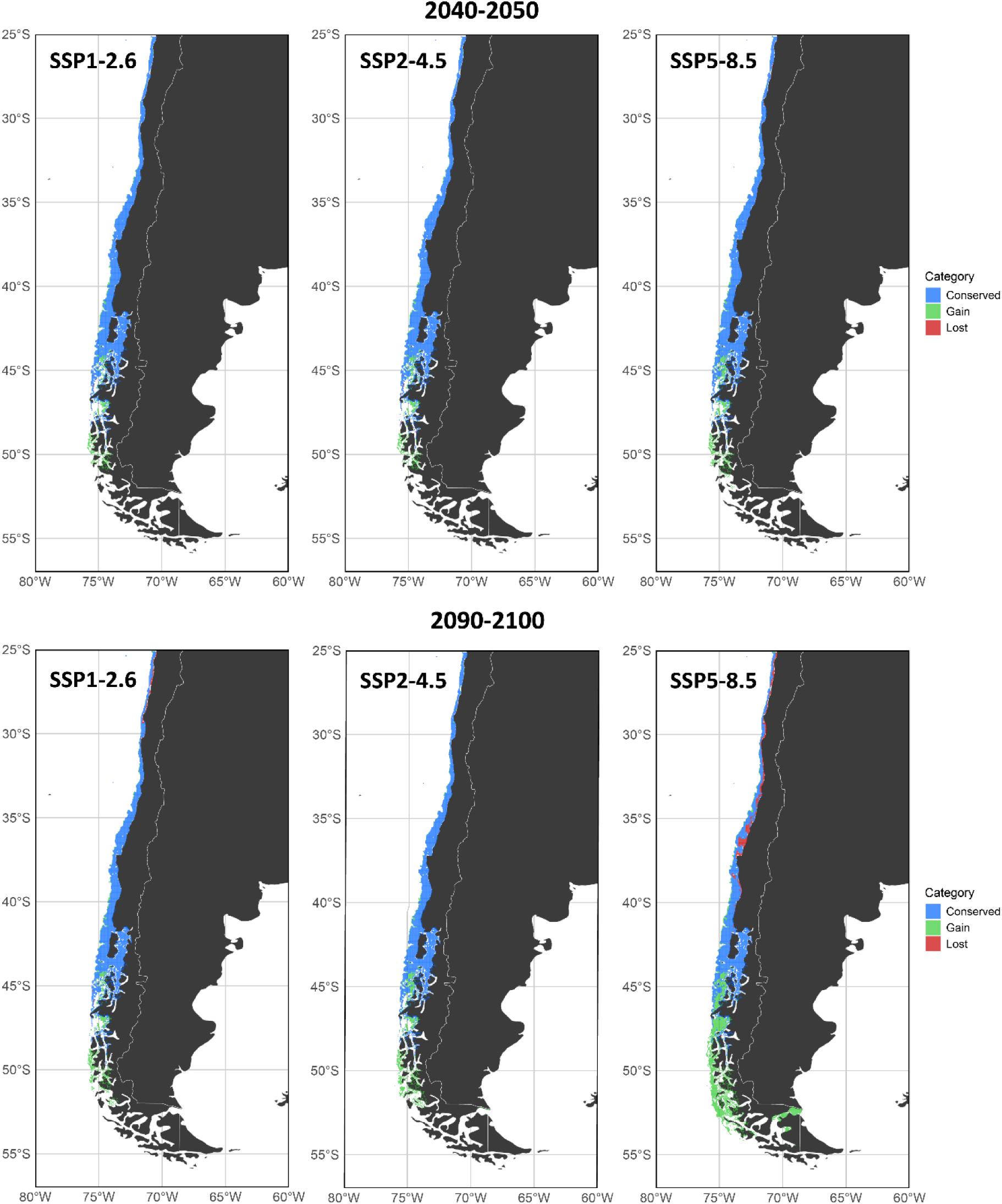
Suitable habitat of *Dipturus chilensis* across the study area. The figure contrasts the present distribution with projections under future climate scenarios (SSP1-2.6, SSP2-4.5, and SSP5-8.5) for 2040–2050 and 2090–2100. Areas of habitat conservation are shown in blue, losses in red, and gains in green.

Across all scenarios, the total suitable habitat for *D. chilensis* was projected to increase from present conditions by the end of the century (**Fig. 3**). The present-day estimate averaged approximately 132,000 km². Projections for 2040–2050 indicated moderate increases, ranging from ∼147,400 km² under SSP 2-4.5 and SSP1-2.6 to ∼151,800 km² under SSP 5-8.5. By 2090–2100, habitat expansions became more pronounced, with estimates reaching ∼148,400 km² under SSP 2-6.0, ∼155,600 km² under SSP 2-4.5, and up to ∼173,400 km² under SSP 5-8.5. Although the magnitude of increase varied by emission pathway, all scenarios consistently predicted a larger area of suitable habitat compared to the present, with the most substantial expansion occurring under the highest-emission scenario (**Table S3**).

**Figure 3.**
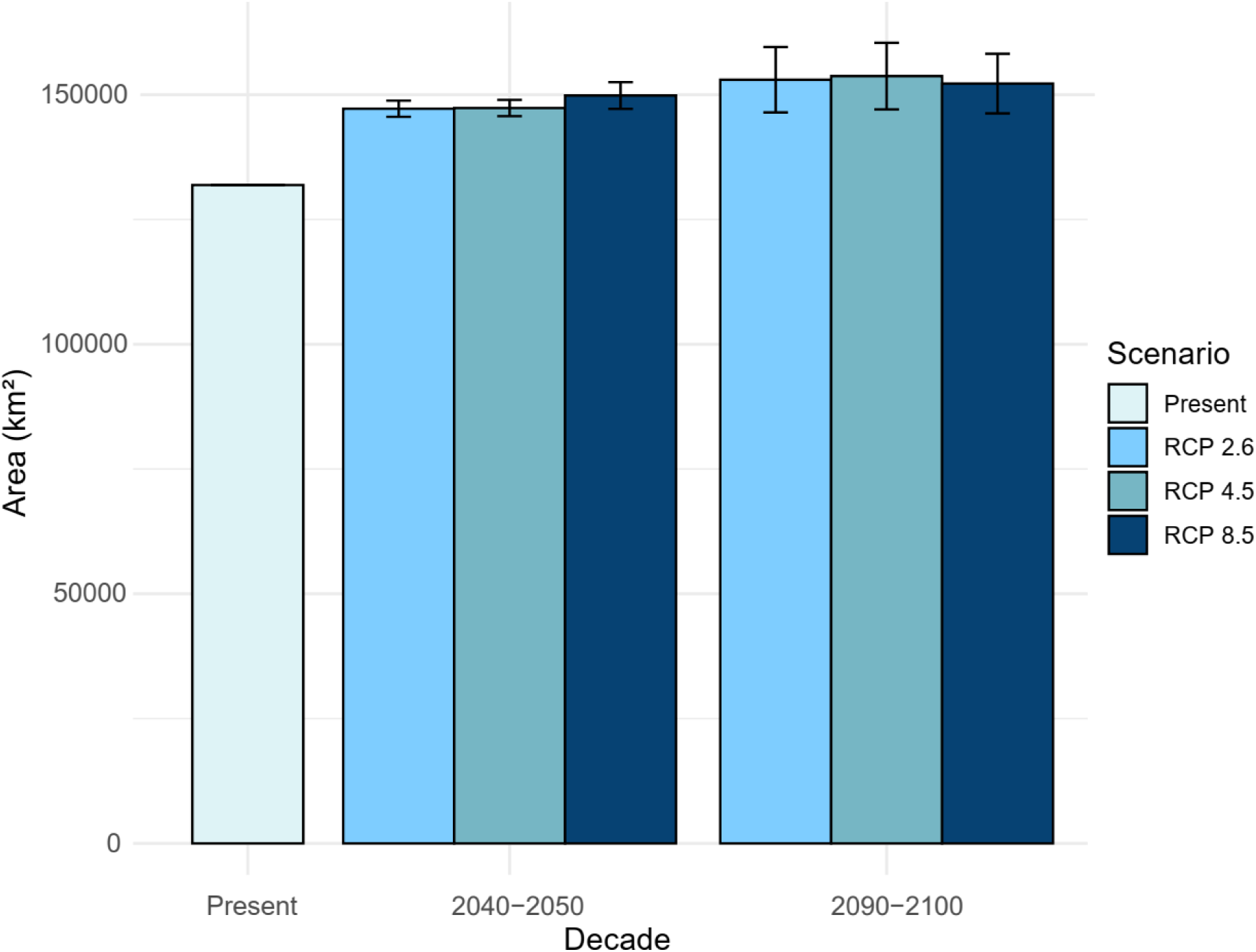
Suitable habitat area for *Dipturus chilensis* under current and future climate scenarios (SSP1-2.6, SSP2-4.5 and SSP5-8.5) for present, 2040-2050 and 2090-2100. Bars show total estimated area (millions of km²), with the lightest blue for present, and darker shades for future climatic scenarios. Error bars represent standard deviation from ten model replicates, indicating variability. Note the overlap between the present error bar and its bar edge.

## 4. Discussion

Our models demonstrated robust predictive performance. Maximum temperature at depth and minimum oxygen at depth were the most influential variables. Projections under different climate scenarios suggest a net expansion of suitable habitat for *D. chilensis* towards southern Chile, particularly in the Channels and Fjords of Southern Chile ecoregion, alongside localized losses in the Central Chile and Araucanian ecoregions of its current range. These results highlight a climate-driven habitat redistribution that could reshape the ecology and conservation priorities of this vulnerable skate in the southeastern Pacific.

The observed poleward expansion in *D. chilensis* is consistent with broad thermal responses reported for marine fishes and elasmobranchs in response to ocean warming (Nye et al., 2009; Dulvy et al., 2014). However, this pattern is not universal. While some benthic skates do not show major geographic shifts under moderate warming, likely because the new conditions remain within their fundamental niche (Coelho et al., 2022), other global models project redistributions exceeding 1,000 km for marine megafauna (Womersley et al., 2024). The particular vulnerability of egg-laying skates is underscored by experimental evidence, which shows that warming and acidification impair embryonic development and survival (Di Santo, 2015; Ripley et al., 2021). Moreover, the compounding threat of overfishing can accelerate extinction risk, as demonstrated for endemic rays in the southwest Indian Ocean (Pollom et al., 2024). Therefore, although the projected habitat expansion in the Channels and Fjords of Southern Chile ecoregion may offer a climatic refuge for *D. chilensis*, its persistence is likely threatened by its physiological sensitivity and the potential for synergistic impacts with other anthropogenic stressors. Notably, fishing pressure has already caused local extirpation of other large-bodied *Dipturus* skates from European waters (Brander, 1981; Rogers and Ellis, 2000; Garbett et al., 2023; Loca et al., 2005.), likely interacting synergistically with environmental stressors and compromising the species long term persistence.

The projected habitat redistribution also has important ecological implications. Expansion into sub-Antarctic ecoregions (i.e., the Channels and Fjords of Southern Chile ecoregion) under temperature-increase models may create refugia characterized by cooler temperatures and higher oxygen availability, offering conditions conducive to persistence. Conversely, habitat contraction in Central Chile and Araucanian ecoregions could fragment population connectivity, generating a spatial pattern of local extinction and distal colonization. Such changes are likely to alter benthic community dynamics by reshaping prey availability, modifying competitive interactions, and exposing *D. chilensis* to novel ecological assemblages (Simpfendorfer and Kyne, 2009; Schwieterman et al., 2019). Furthermore, the redistribution of skates and other benthic predators has been shown to restructure ecosystems under the combined pressures of climate change and fishing (Sguotti et al., 2016), a dynamic that may also emerge in the southeastern Pacific.

Despite the predicted expansion of suitable habitat, our results do not imply a reduced extinction risk for *D. chilensis*. The species remains listed as Endangered by the IUCN (Dulvy et al., 2021), due to its low fecundity, slow growth, late maturity, and history of overexploitation (Quiroz et al., 2011; Wiff et al., 2018; Concha et al., 2012). Global assessments confirm that large-bodied, coastal elasmobranchs are disproportionately vulnerable to the combined threats of fishing and climate change (Dulvy et al., 2014). Regional analyses from the Eastern Tropical Pacific further highlight elevated climate vulnerability in batoids and deep-water chondrichthyans (Cerutti-Pereyra et al., 2024), reinforcing concerns for temperate demersal skates. Furthermore, as the species shifts polewards, fishing effort may follow into potentially under-regulated areas. Consequently, conservation planning must anticipate not only habitat gains but also the socio-ecological feedback of this redistribution. Establishing or expanding Marine Protected Areas in southern Chile, integrated with adaptive fisheries management, could be crucial for the conservation of *D. chilensis*. Unlike highly migratory taxa, many mid-and large-bodied skates, including *Dipturus* spp. exhibit spatially restricted movement and area-focused habitat uses (Wearmouth and Sims, 2009; Siskey et al., 2019; Ahumada et al., 2025), with distinct phylopatric patterns being reported for a handful of species (Flowers et al., 2016; Hillinger et al., 2025). Consequently, strategically located Marine Protected Areas in southern Chile could form a valuable component of a conservation strategy. To account for the impacts of climate change, however, these static protections must be paired with adaptive fisheries management, and a regional approach that can adjust to shifts in distribution and population connectivity (Garbett et al., 2021; Bisch et al., 2022). Although our occurrence dataset shows fewer records in the Channels and Fjords of Southern Chile ecoregion, this scarcity might reflect limited scientific sampling and reporting in a remote, logistically challenging area rather than true biological absence. We addressed this imbalance through spatial thinning (11 km) and a KDE-derived sampling-bias layer that was supplied to Maxent, thereby down-weighting the influence of oversampled central regions. While the robustness of our projections is supported by the robust discrimination ability of our model (empirical AUC = 0.94 vs. null models AUC = 0.51, **Table S2)**, predictions in data-poor areas inherently carry higher uncertainty. Autoecological studies in southern Chile (e.g., Ahumada et al., 2025) will be essential to validate and refine these predictions.

Future research could integrate species distribution models with movement data, such as those provided by remote sensing studies (e.g., Ahumada et al., 2025), to validate projected habitat shifts. Model realism would also benefit from incorporating biotic interactions and direct fishing pressure alongside abiotic variables, while also addressing potential biases from uneven occurrence records (Robinson et al., 2017). Integrated models that account for the interactive effects of climate and fisheries on population dynamics are particularly needed (Dulvy et al., 2003; Frisk et al., 2005). Extending this approach to sympatric skates and sharks would help identify community-level redistribution patterns and anticipate ecosystem consequences. Ultimately, developing adaptive management strategies that incorporate climate projections into conservation policy is essential to enhance the resilience of fisheries and protect vulnerable elasmobranchs in the southeastern Pacific.

Our study projects a net poleward redistribution of *D. chilensis*, with a climate-driven expansion of suitable habitat into the Channels and Fjords of Southern Chile ecoregion. However, these potential gains do not automatically translate into viable populations, and their persistence may depend on proactive conservation measures and adaptive fisheries management. We therefore emphasize that climate-informed governance is urgently needed to ensure the long-term persistence of this emblematic skate in the southeastern Pacific.

## Acknowledgements

This project was partially funded by the Fondo Nacional de Desarrollo Científico y Tecnológico [ANID/FONDECYT 3230610, and ANID/FONDECYT 1251475], ANID-CENTROS REGIONALES (CLAP R20F0008), and CICYT BIP 40069515-0 . We thank the Instituto de Fomento Pesquero (IFOP) for providing part of the data necessary for the analyses. We also acknowledge the Global Biodiversity Information Facility (GBIF) and the data publishers for making biodiversity occurrence data openly available. We thank the reviewers of this study and the editor for their constructive comments that helped improve the manuscript.

## Author contributions

**Jaime A, Villafaña:** Conceptualization, Writing – original draft, Writing – review and editing. **Diego Almendras:** Conceptualization, Writing – original draft, Writing – review and editing. **Daniel González-Aragón:** Formal analysis, Methodology, Software, Visualization, Writing – original draft. **Felipe I. Torres:** Formal analysis, Methodology, Software, Visualization, Writing – original draft. **Francisco Concha:** Supervision, Writing – review and editing. **Ana Guzman-Castellanos:** Supervision, Writing – review and editing. **Ignacio Contreras:** Data curation, supervision, Writing – review and editing. **Karina E. Buldrini:** Writing – review and editing. **Pablo Oyanadel-Urbina:** Writing – review and editing. **Carolina A. Sandoval:** Writing – review and editing. **Benjamín Miranda:** Data curation, **Gabriel Mazo:** Data curation, **Felipe Cárdenas:** Data curation, **Mathias Valdivia:** Data curation, **German Pequeño:** Supervision, Writing – review and editing. **Carlos Lara:** Supervision, Writing – review and editing. **Marcelo M. Rivadeneira:** Supervision, Writing – review and editing.

## Ethics approval consent to participate

No approval of research ethics committees was required to accomplish the goals of this study

---

**Figure S1.**
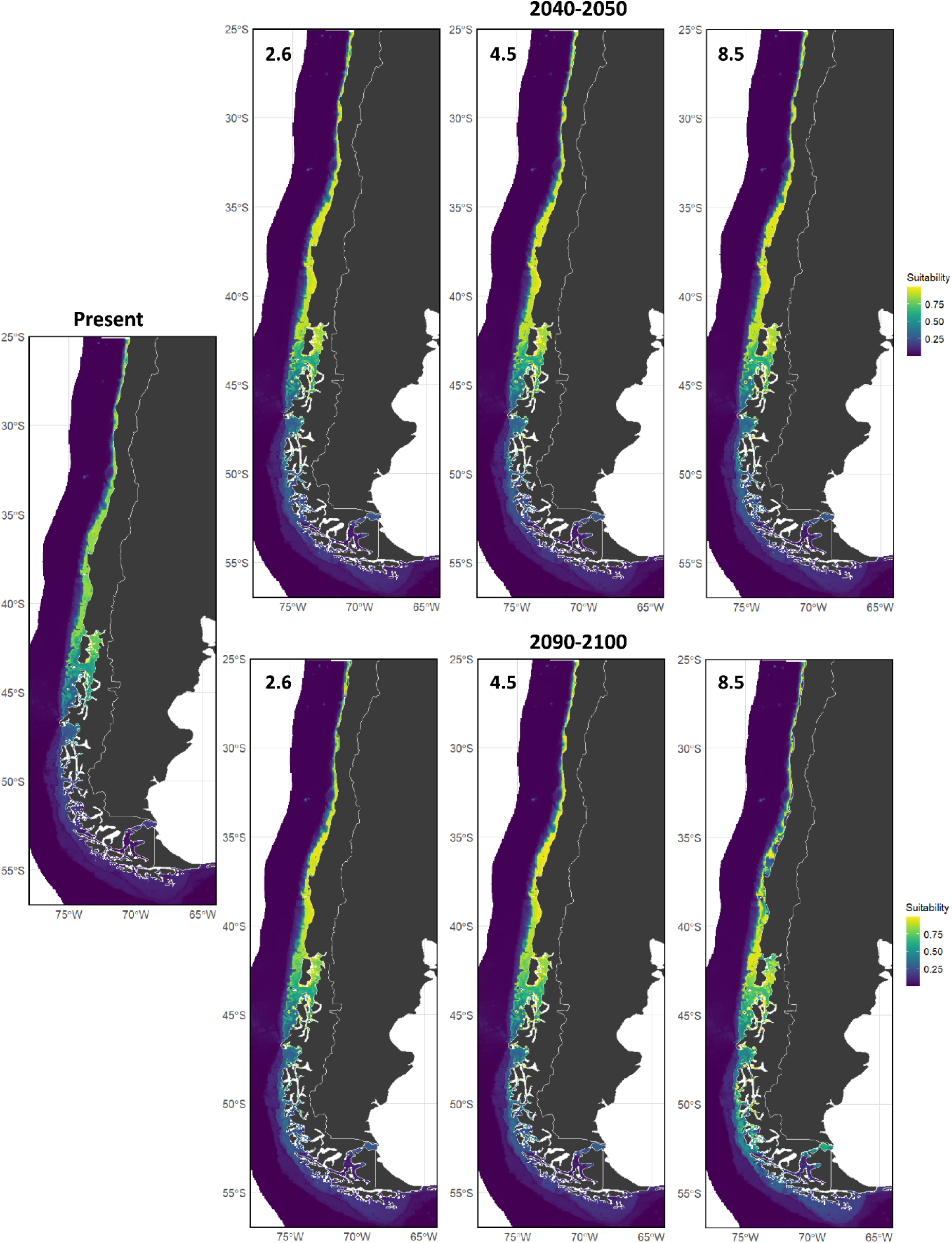
Suitable habitat area for *Dipturus chilensis* under current and future climate scenarios (SSP1-2.6, SSP2-4.5 and SSP5-8.5) for 2040-2050 and 2090-2100.

**Table S1.** Georeferenced occurrences of *Dipturus chilensis* along the coast of Chile, with the source for each record indicated.

**Table S2.** Empirical versus null model performance, including Wilcoxon test results assessing whether empirical performance exceeds random expectation.

**Table S2.** Changes in suitable habitat area (km²) of *Dipturus chilensis* under different future climate scenarios (SSP2-6.0, SSP4-5.0, and SSP8-5.0) for 2050 and 2100, showing conserved, gained, and lost areas relative to the present distribution.

